## Supplementary Material for "Neurofeedback training can modulate task-relevant memory replay rate in rats"

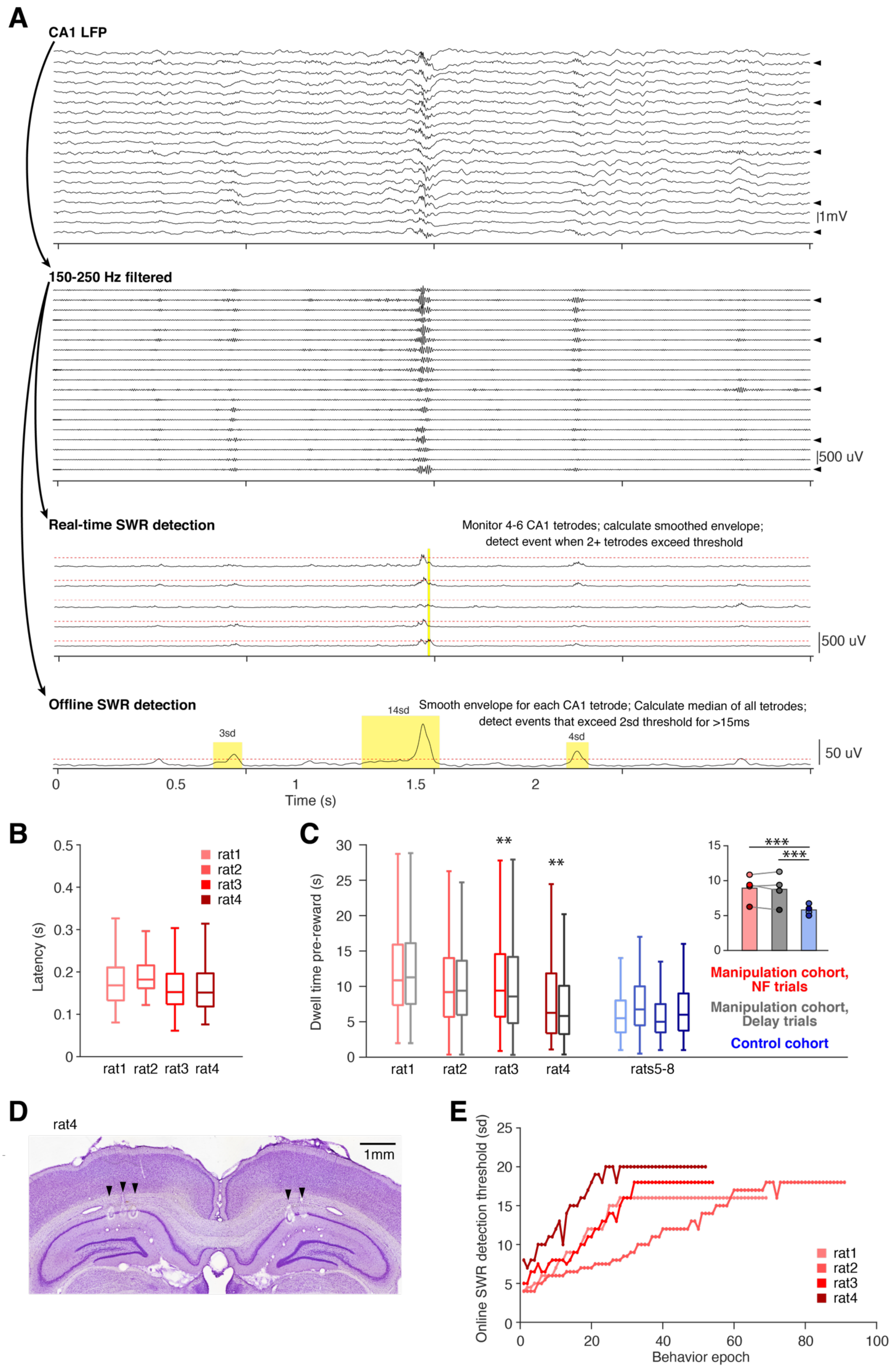

A
CA1 LFP
1mV
150-250 Hz filtered
500 uV
Real-time SWR detection
Monitor 4-6 CA1 tetrodes; calculate smoothed envelope;
detect event when 2+ tetrodes exceed threshold
500 uV
Offline SWR detection
Smooth envelope for each CA1 tetrode; Calculate median of all tetrodes;
detect events that exceed 2sd threshold for >15ms
3sd
14sd
4sd
50 uV
Time (s)
B
Latency (s)
rat1
rat2
rat3
rat4
C
Dwell time pre-reward (s)
rats5-8
Manipulation cohort, NF trials
Manipulation cohort, Delay trials
Control cohort
D
rat4
1mm
E
Online SWR detection threshold (sd)
Behavior epoch

**Supplementary Figure 1. A neurofeedback approach to reinforcing SWRs, related to Figure 1.**
(A) Schematic of the online and offline SWR detection strategies used. For both detection methods, raw LFP from tetrodes located in CA1 cell layer are filtered for ripple band power (100-400 Hz bandpass filter for online detection; 150-250 Hz bandpass filter for offline detection). For online detection, 4-6 tetrodes are chosen each day (marked with arrowheads). During a period of movement just prior to the start of the behavioral epoch, the envelope of the ripple filtered signal is calculated for the chosen tetrodes, the mean and standard deviation (sd) of the envelope are calculated, and are fixed for the subsequent behavioral epoch. A threshold is set as a certain number of sd above the mean and events which cross this threshold on at least two chosen tetrodes simultaneously are detected as suprathreshold SWRs. In contrast, the offline detection strategy calculates the envelope of the ripple-filtered trace for all CA1 cell layer tetrodes and calculates a "consensus trace" as the median across tetrodes (see Methods). The mean and sd of this consensus trace is calculated and events which exceed 2 sd above the mean for at least 15 ms are considered SWR events.
(B) Feedback latency to reward delivery for the manipulation cohort. Feedback latency is calculated as the time between the start of the offline-detected SWR event and the tone/reward delivery. Note that this latency includes both the time for the event to reach the online detection threshold on multiple tetrodes as well as the fixed delay added to reduce the chance of feedback delivery interrupting the ongoing SWR event (100, 50, 75, and 75 ms for each subject, respectively). n = 3948, 2155, 3880, and 4894 suprathreshold events per subject, respectively.
(C) Time spent at the center ports pre-reward for subjects of each cohort. Manipulation cohort n = 1892, 684, 1157, and 1602 NF trials and 2022, 640, 1201, and 1552 delay trials; control cohort n = 2490, 2629, 2027, and 3021 trials, respectively. Manipulation cohort ranksum comparisons between neurofeedback (NF) and delay trials: $p = 0.228$, 0.690, 0.005, and 0.002, respectively. Inset: Groupwise comparisons. Manipulation cohort NF trials vs control cohort trials: $p = 2.936 \times 10^{-9}$; manipulation cohort delay trials vs control cohort trials: $p = 2.178 \times 10^{-5}$.
(D) Representative histology example from rat4 showing lesions (arrowheads) at locations where tetrode tips were located in dorsal CA1 cell layer.
(E) Maximum online detection threshold for SWRs for each behavioral epoch of neurofeedback training for subjects in the manipulation cohort.

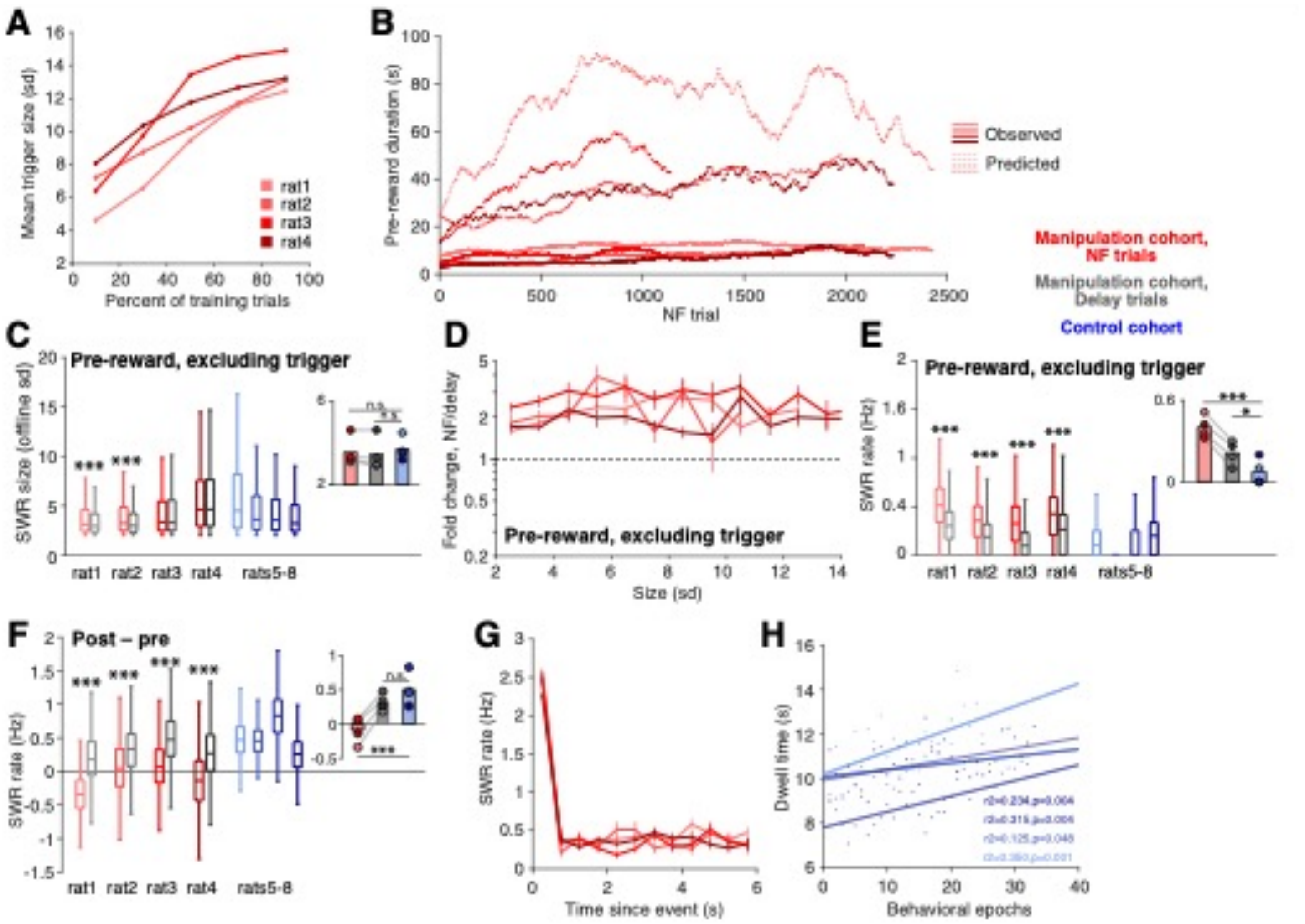


**Supplementary Figure 2. Neurofeedback training modulates SWR rate, related to Figure 2.**

(A) Mean trigger event size (sd) for neurofeedback trials over the early portion neurofeedback training (while increasing the detection threshold). Trials included are those from the second half of each behavioral epoch to ensure that the online detection threshold had been raised to its maximum value (see Methods).

(B) Actual duration pre-reward (solid lines) compared to predicted duration (dashed lines) based on the occurrence rate of large-amplitude SWRs prior to and early in neurofeedback training. Traces have been smoothed with a 200-trial window (see Methods).

(C) SWR size during the pre-reward period at the center ports. All suprathreshold events for both trial types within the manipulation cohort are excluded. Manipulation cohort n = 13209, 3060, 5369, and 7165 NF SWRs and 8716, 1618, 2535, and 3855 delay SWRs; control cohort n = 2660, 581, 2192, and 5429 SWRs. Manipulation cohort ranksum comparisons between NF and delay trials: $p = 1.734 \times 10^{-16}$, $7.389 \times 10^{-9}$, 0.921, and 0.921. Inset: Groupwise comparisons. Manipulation cohort NF trials vs control cohort trials: $p = 0.234$; manipulation cohort delay trials vs control cohort trials: $p = 0.184$.

(D) Fold change of occurrence rate of SWRs binned by amplitude at neurofeedback port relative to delay port pre-reward, excluding all suprathreshold events for both trial types.

(E) SWR rate during the pre-reward period calculated with suprathreshold events excluded from both neurofeedback and delay trials in the manipulation cohort. Manipulation cohort n = 1892, 684, 1157, and 1602 NF trials and 2022, 640, 1201, and 1552 delay trials; control cohort n = 2490, 2629, 2027, and 3021 trials. Manipulation cohort ranksum comparisons between NF and delay trials: $p = 9.873 \times 10^{-136}$, $2.562 \times 10^{-34}$, $4.017 \times 10^{-91}$, and $7.171 \times 10^{-49}$. Inset: Groupwise comparisons. Manipulation cohort NF trials vs control cohort trials: $p = 5.574 \times 10^{-8}$; manipulation cohort delay trials vs control cohort trials: $p = 0.040$.

(F) The difference between SWR rate during the post-reward period and SWR rate during the pre-reward period. Manipulation cohort n = 1892, 684, 1157, and 1602 NF trials and 2022, 640, 1201, and 1552 delay trials; control cohort n = 2490, 2629, 2027, and 3021 trials. Manipulation cohort ranksum comparisons between NF and delay

trials: $p = 0$, $2.132 \times 10^{-36}$, $4.528 \times 10^{-117}$, and $3.821 \times 10^{-139}$. Inset: Groupwise comparisons. Manipulation cohort NF trials vs control cohort trials: $p = 1.352 \times 10^{-5}$; manipulation cohort delay trials vs control cohort trials: $p = 0.139$.
(G) SWR rate in 0.5 s bins calculated following the detection of suprathreshold events during delay trials, pre-reward, with at least 2 s remaining prior to reward delivery, shows no extended suppression of SWR rate following suprathreshold events.
(H) For the control cohort, the amount of time spent post reward delivery at the center well tends to increase as subjects gain more experience with the task.
For C, E, and F, all ranksum p-values are corrected using the Benjamini-Hochberg method and all groupwise comparisons are performed using linear mixed effects models (see Methods)

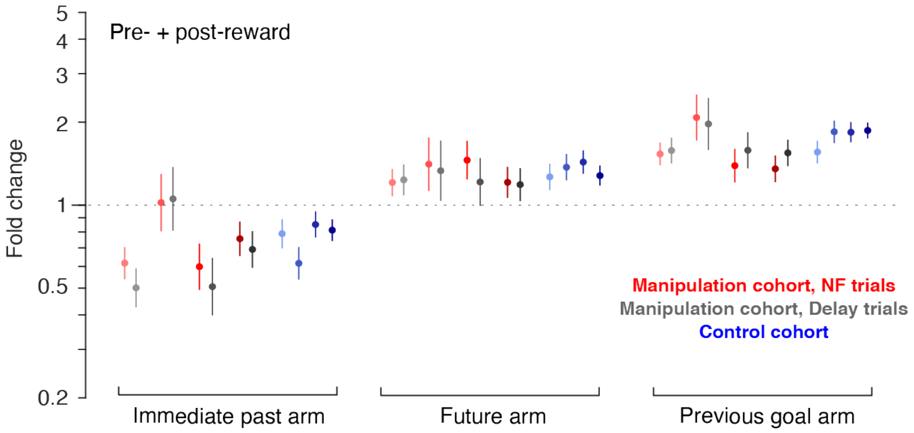


**Supplementary Figure 3. Replay content is consistent when considering all replay at the center ports, related to Figure 4.**
GLM quantifying modulation of replay rate by arm category considering replay events during both the pre- and post-reward periods at the center ports. Manipulation cohort n = 1661, 392, 866, and 1281 neurofeedback (NF) trials and 1705, 367, 894, and 1213 delay trials; control cohort n = 1458, 1636, 1464, and 2181 trials, respectively.

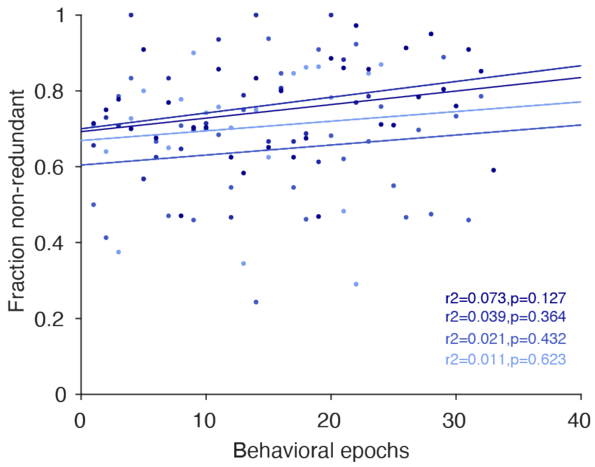


**Supplementary Figure 4. Search efficiency increases with experience, related to Figure 5**.
For the control cohort, search efficiency tends to increase over behavioral epochs.
